# Rapid ageing and species identification of natural mosquitoes for malaria surveillance

**DOI:** 10.1101/2020.06.11.144253

**Authors:** Doreen J. Siria, Roger Sanou, Joshua Mitton, Emmanuel P. Mwanga, Abdoulaye Niang, Issiaka Sare, Paul C.D. Johnson, Geraldine Foster, Adrien M.G. Belem, Klaas Wynne, Roderick Murray-Smith, Heather M. Ferguson, Mario González-Jiménez, Simon A. Babayan, Abdoulaye Diabaté, Fredros O. Okumu, Francesco Baldini

## Abstract

The malaria parasite, which is transmitted by several *Anopheles* mosquito species, requires more time to reach its human-transmissible stage than the average lifespan of a mosquito. Monitoring the species-specific age structure of mosquito populations is critical to evaluating the impact of vector control interventions on malaria risk. We developed a rapid, cost-effective surveillance method based on deep learning of mid-infrared spectra of mosquitoes’ cuticle that simultaneously identifies the species and the age of three main malaria vectors, in natural populations. Using over 40,000 ecologically and genetically diverse females, we could speciate and age grade *An. gambiae, An. arabiensis*, and *An. coluzzii* with up to 95% accuracy. Further, our model learned the age of new populations with minimal sampling effort and detected the impact of control interventions on simulated mosquito populations, measured as a shift in their age structures. We anticipate our method to be applied to other arthropod vector-borne diseases.

---

Malaria presents a paradox: its transmission depends on mosquito vectors that have a shorter mean lifespan than the malaria parasite requires for its own development^1^. Consequently, its persistence depends on the small proportion of mosquitoes that live long enough to transmit malaria sporozoites to a suitable mammalian host. This causes small changes in mosquito longevity to have a big impact on malaria transmission^2^, which explains why malaria control has focussed on interventions that primarily target adult mosquito survival^3^. Examples include insecticidal nets which have substantially reduced the incidence of malaria^4^, but may now be threatened by insecticide resistance^5^. A further consequence of those mosquito/malaria life cycle dynamics is that accurate and reliable assessment of mosquito age structure is crucial for monitoring the impact of vector control interventions. However, current mosquito age grading methods typically rely on 60-year-old techniques based on ovary dissections^6,7^ that are slow, labour-intensive and imprecise, and which vary between mosquito species^8^. Many alternatives have been investigated with uneven success^9–14^. Because malaria is transmitted by multiple, often morphologically indistinguishable mosquito species that differ in longevity, behaviours, and vectorial capacity^15,16^, a method that simultaneously estimates vector species and age without relying on time-consuming techniques and expensive reagents would be of great value.

Like all arthropods, mosquitoes have a cuticle whose chemical composition differs between species and changes with age^8^, which infrared spectroscopy can detect by quantifying how the mosquito cuticle absorbs light^13,17,18^. Early work on infrared spectroscopy for mosquito analysis was restricted to the near-infrared spectrum (10,000–4,000 cm^−1^)^13,18,19^.

While Near-Infrared Spectroscopy (NIRS) can distinguish species and age groups in laboratory settings, it has not been shown to predict the age of mosquitoes in natural environments^20^, where natural genetic and ecological variability are expected to affect how mosquito cuticle develops over time and between populations even of the same species. Mid-Infrared Spectroscopy (MIRS, 4,000 - 400 cm^−1^) is an alternative technology that, unlike NIRS, measures discrete fundamental vibrations of biomolecules, allowing more information to be extracted from biological samples (such as on protein conformation)^21,22^ and therefore detecting more subtle changes among species or mosquitoes of different ages. Recently, we demonstrated that MIRS can accurately predict the species and age structure of African malaria vectors under controlled environmental conditions^23^ although its applicability to ecologically and genetically variable wild mosquitoes was not tested.

In this study, we developed a MIRS approach to predict species and age of natural populations of three major African malaria vectors. We used a deep learning MIRS (DL-MIRS) model based on geographically and ecologically distinct female mosquitoes in East and West Africa to re-construct the age structure of natural mosquito populations and detect their changes pre- and post-simulated vector control interventions. These results demonstrate how this low-cost, artificial intelligence-based approach can quantify previously immeasurable impacts of interventions on natural vector populations, and constitute a new surveillance tool in the fight against malaria.

## Results

### Ecologically and genetically variable dataset built for natural mosquito population surveillance

We created a dataset of mosquito MIR spectra from diverse genetic backgrounds and reared both in diverse laboratories and in ecologically-realistic semi-field systems in East and West Africa to capture laboratory (LV), genetic (GV), and environmental variation (EV) (Fig. 1). This dataset comprised 41,151 female mosquitoes of three *An. gambiae s.l*. group species, *An. gambiae, An. arabiensis*, and *An. coluzzii* from diverse origins (Supplementary Table 1): the LV subset comprised spectra of mosquitoes from three different laboratories from the UK, Tanzania, and Burkina Faso; GV and EV included spectra from adult mosquitoes collected from the field (as eggs or larvae) or derived from laboratory colonies in Tanzania and Burkina Faso, and reared in laboratory or semi-field systems, respectively (Supplementary Table 2). Using semi-field enclosures allowed us to rear mosquitoes under natural environmental variability while controlling their age. To account for the variation within and between these diverse populations, we trained convolutional neural networks (CNNs) on MIR spectra from this dataset to simultaneously predict both age and species using data collected from dried mosquitoes by an attenuated total reflectance (ATR) Fourier-transform infrared (FTIR) spectrometer.

**Fig. 1.**
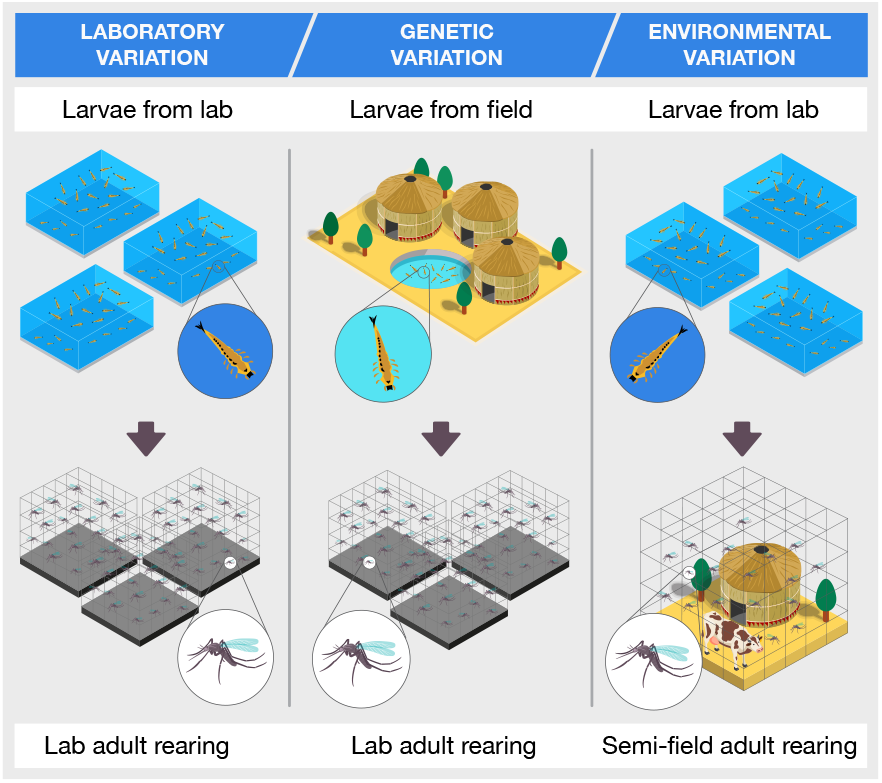
Experimental setup for capturing variation in MIRS caused by laboratory of origin, individual genetic differences, and natural environment. To disentangle genetic and environmental effects mosquitoes were obtained from either laboratory-bred colonies or from genetically heterogeneous wild larvae; half of the laboratory larvae were then reared and allowed to develop through adult stage in semi-field conditions, which offer ecologically-realistic conditions while still allowing control of mosquito age.

### DL-MIRS is sensitive to mosquito cuticle biochemical signature and accurately predicts mosquito age and species

First, we used unsupervised clustering of all MIRS data regardless of their origin using Uniform Manifold Approximation and Projection (UMAP)^25^ revealed signatures of both geographic origin and development environment in all three *Anopheles* species (Fig. 2a,b).

**Fig. 2.**
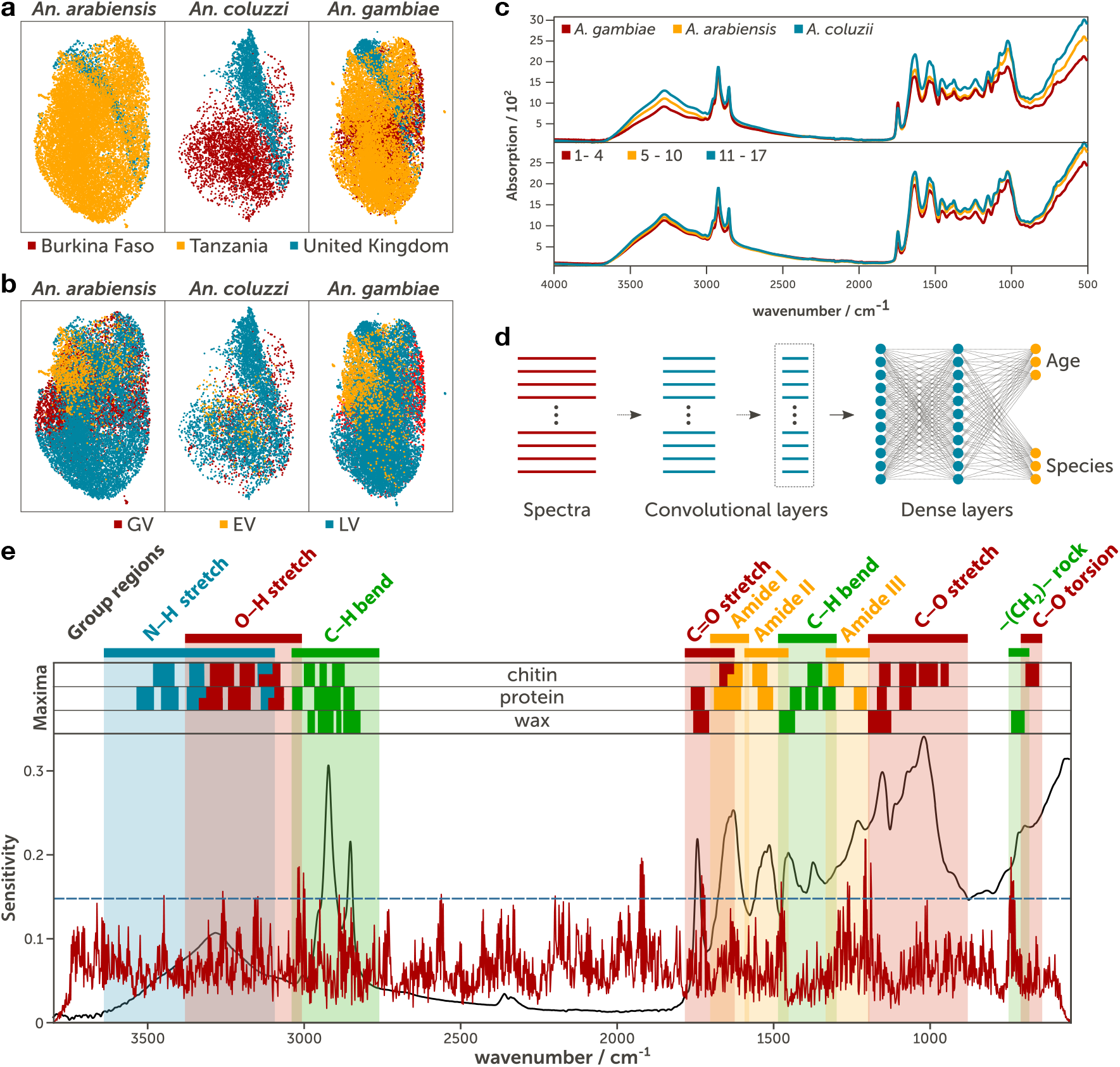
Variation in MIRS, machine learning model architecture, sensitivity of the trained model. We collected the MIRS of 41,151 female mosquitoes belonging to three species from diverse laboratories, genetic backgrounds, and environments and three age classes spanning 1—17 days post pupal emergence. **a, b**, Unsupervised clustering of MIRS measurements using Uniform Manifold Approximation and Projection of MIRS in two dimensional space (plot axes) from *An. arabiensis, An. coluzzii*, and *An. gambiae* coloured according to site of origin (a) and source of variation (b). **c**, Representative variation of mid-infrared absorption spectra of *An. arabiensis, An. coluzzii*, and *An. gambiae* and of three age classes. **d**, Schematic representation of the deep convolutional neural network that takes MIRS inputs and outputs mosquito age and species. The input layer (wavenumbers) is fed through five 1-dimensional convolutional layers, comprising of 16 filters each (convolutional layers region), followed by a dense layer of 500 features and age and species output layers (dense layers) that were used to make predictions. **e**, Average model sensitivity (red line) to different wavenumbers of all 41,151 input spectra and comparison with the features of a representative absorption spectrum of a mosquito (grey line). The dashed blue line represents 95% confidence in the sensitivity values^24^. The coloured stripes show the regions associated with the particular vibration of a functional chemical group. The upper part (maxima) displays the intervals of wavenumbers in which the maximum of the absorption peaks of each vibration appear for each of the three most abundant components in the cuticle of a mosquito^23^. Here, the vibration of the same bonds appear in different wavenumbers depending on which cuticular component they belong to (chitin, protein or wax), which modifies the shape of the peaks.

We then used a supervised learning approach to disentangle patterns associated with mosquito origin from any variation specific to their species and age within the MIRS data. Mosquitoes MIRS were then grouped by species and three age classes (days after pupal emergence) corresponding to their potential to be infected and transmit malaria (Fig. 2c): younger non-infected (1–4 days old), potentially infected but not infectious (5–10 days old) and old enough to be infectious (≥11 days old). Each MIR spectrum was used as an input of a deep CNN and the final optimised architecture included five 1-D convolutional layers followed by dense and prediction layers (Fig. 2d). Sensitivity analysis of the input spectra showed that the model was particularly sensitive to wavenumbers corresponding to C–O, O–H, and C=H stretches, C–H bends, amide III, and −CH_2_–rock structures within the mosquito cuticle (Fig. 2e and Supplementary Fig. 1a). First, we trained and tested the DL-MIRS model to account for laboratory variation with the LV dataset alone, achieving which 84% and 93% accuracy in classifying age and species, respectively (Fig. 3a,b). Then, training and testing were performed on both LV and GV datasets to capture genetic variation, and prediction of age and species of wild-derived adult females achieved 89% and 95% accuracy, respectively (Fig. 3c,d). We then tested how well models trained on both LV and GV datasets could predict age and species of mosquitoes that developed in ecologically-realistic semi-field environmental conditions (EV). However, these models performed no better than chance (Fig. 3e,f) regardless of whether we preselected input wavenumbers as described previously^23^ (Supplementary Table 3 and Fig. 1b). We hypothesised that the failure of these models to predict the age and species of “natural” mosquitoes was due to the variation in MIRS profiles being mostly driven by non-genetic factors and therefore that DL-MIRS requires training on some local ecological variation.

**Fig. 3.**
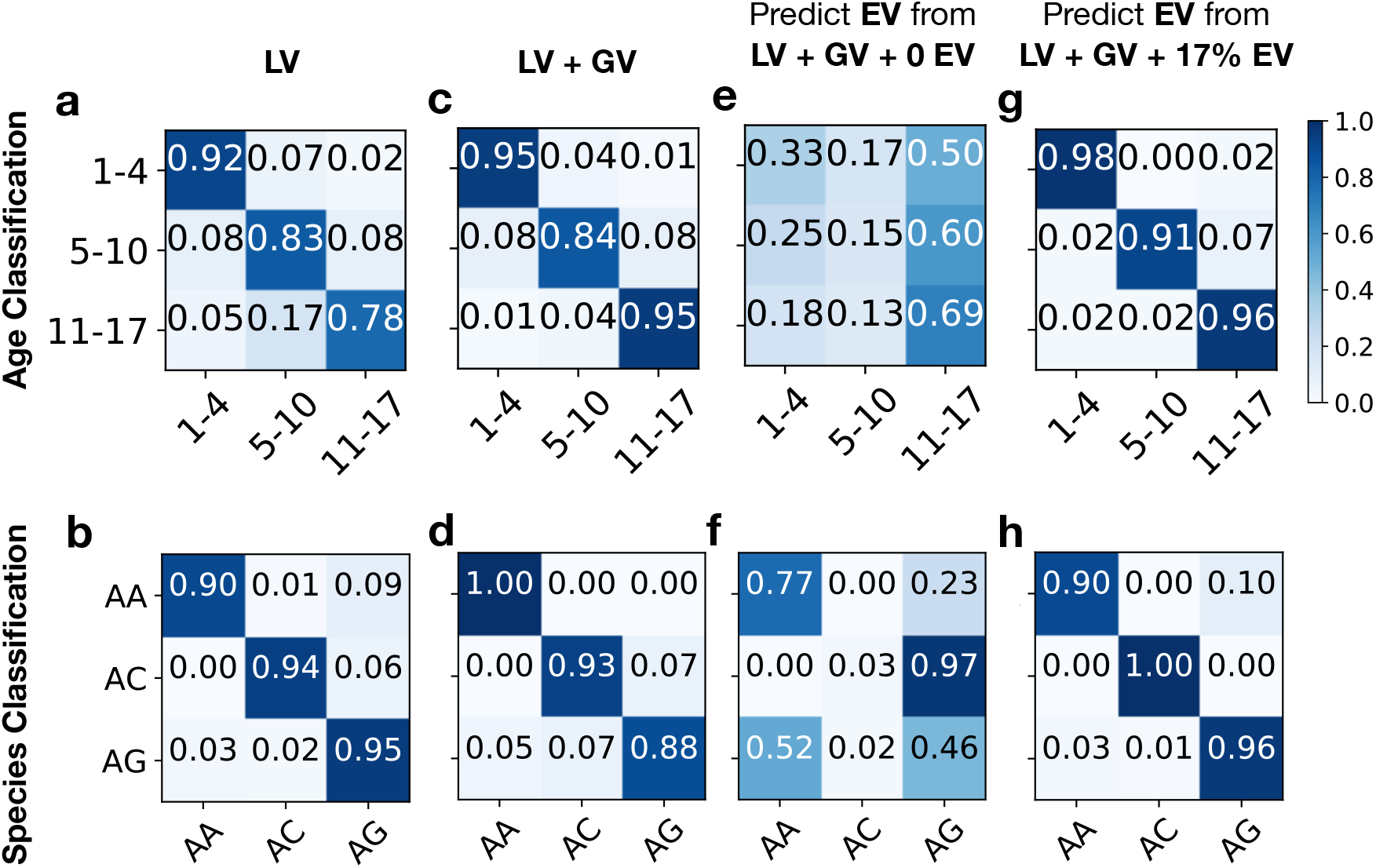
Confusion matrices of model prediction accuracies. DL-MIRS was trained using different combinations of mosquitoes from either laboratory larvae reared in the lab (LV, laboratory variation), larvae from the field reared in the lab (GV, genetic variation), or laboratory larvae reared in semi-field (EV, environmental variation). **a—d**, The models were trained and tested on mosquitoes of the same origins (a, b LV alone; or c, d LV+GV) for their ability to classify a random stratified hold-out test set into the correct age class (a, c) or species (b, d). **e**, **f**, We then trained models on LV+GV mosquitoes, and tested their accuracy in correctly identifying EV mosquito age (e) and species (f). **g**, **h**,To improve model generalisation from lab to field-reared mosquitoes, we used transfer learning by freezing the convolutional layers and re-training only the dense layers with a small proportion of EV mosquitoes (here, 1452 examples), resulting in more accurate identification of mosquito age (g) and species (h).

We therefore performed a series of additional analyses to improve the generalisability of this approach (Supplementary Table S3), and found that using transfer learning from a model trained on an LV+GV dataset with only 1294 EV data samples, a relatively small proportion of local specimens, the model achieved 94% accuracy in both species and age class prediction (Fig. 3g,h). To test whether ecological effects were country-specific, we then trained and tested the DL-MIRS with distinct combinations of mosquito origins. Training a CNN including mosquitoes reared in semi-field facilities from one country could not predict age and species of populations from another (Study E2, Fig. 3). Similarly, training a CNN including laboratory-reared mosquitoes from two sites could not predict age and species of those reared at the third (Study E3, Fig. 4), even with pre-selected wavenumbers (Study E4, Fig. 5). Further, reducing training input parameters does not improve generalisation, showing that the CNN algorithm is not overfitting (Supplementary Table 3 (Study E1) and Supplementary Figure 1b). These results suggest that variation in MIRS associated with mosquito age and species can be successfully learned and traits predicted by a CNN from several laboratories, genetic backgrounds and environments, if some spectra (~1200-1300) from the target population are included in training.

**Fig. 4.**
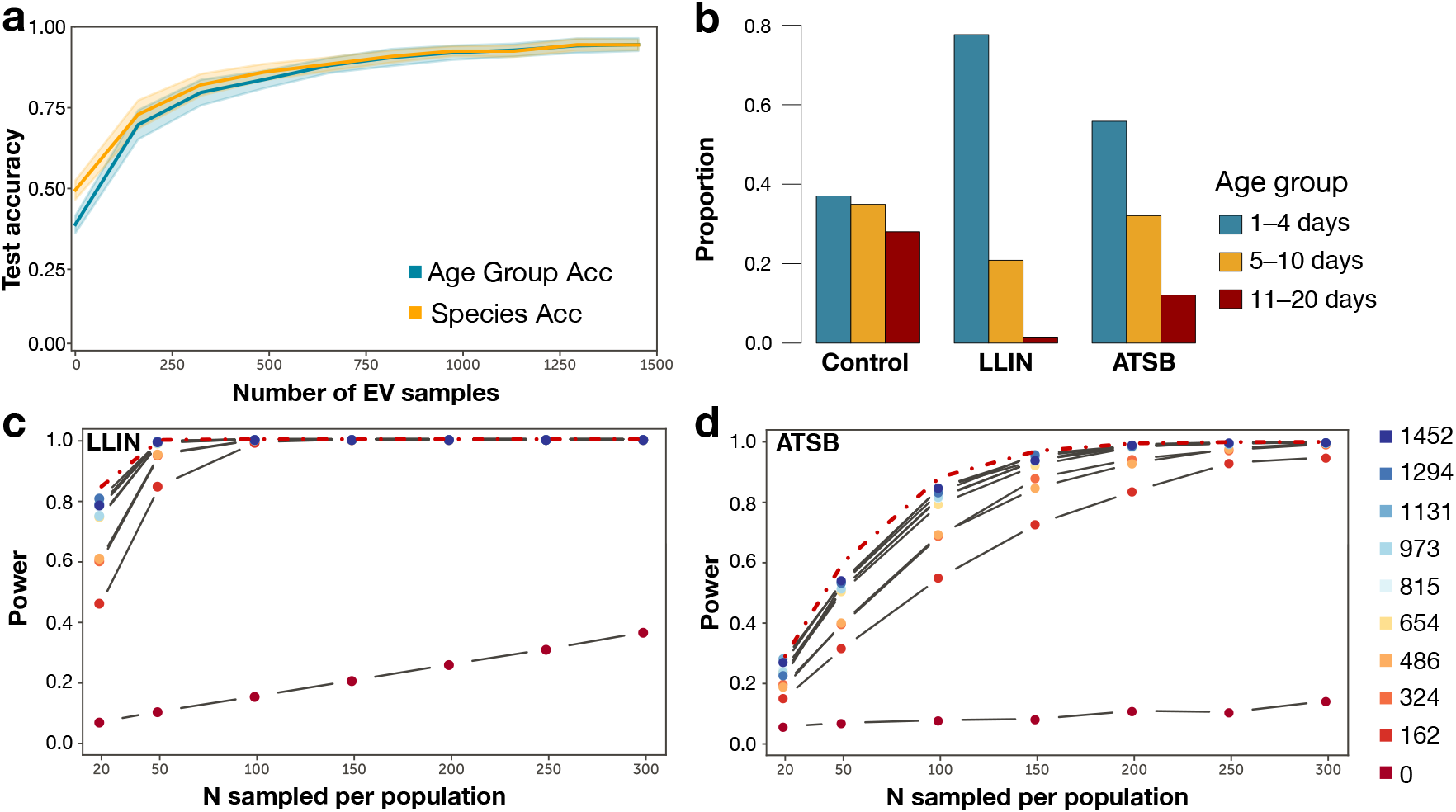
DL-MIRS generalisation and detection of vector control intervention. **a**, Classification accuracy improved from ~50% to 94% for both age group and species with a training set comprising 0 (i.e. effects of increasing sampling of lab-reared mosquitoes only) through 1452 semi-field (EV) mosquitoes used to re-train the transfer learned model. **b**, We used computer simulation to assess the power of DL-MIRS to detect shifts in population age structure in response to each of two vector control interventions, long-lasting insecticide-treated nets (LLIN) and attractive toxic sugar baits (ATSB) relative to a population with no intervention (control). **c**, **d** Power to detect an effect of the vector control intervention was estimated over 10 levels of training set size, with EV mosquitoes ranging from 0 to 1452 and seven sample sizes per population from 20 to 300 (Supplementary Table 4). The dotted red line shows the power that would be achieved with 100% accurate age group classification. The difference between the solid and dotted lines represents the cost in power due to prediction error.

### DL-MIRS transfer learning improves prediction generalisation and minimizes samples required for field re-calibration

To assess the smallest number of local samples required by DL-MIRS to perform adequately while minimising field collection effort, we utilised the features learned by the convolutional layers from the model trained with laboratory data (LV+GV) and transfer learning to train a model for EV data. Here, the model was initially trained end-to-end with LV+GV data, then the convolutional layers (dashed box Fig. 2d) were frozen and the model retrained with EV data only. To test how many samples from the local population were needed, we added incremental proportions of EV data during training. To reduce variation from stochastic processes in model training (Supplementary Fig. 2), model accuracies were averaged. As a result of increasing EV representation in the training set, the prediction accuracy increased rapidly with the inclusion of the target data, exceeding 80% accuracy with 500 examples and exceeding 95% accuracy when over 1000 EV data points were included (Fig. 4a).

Overall, these results suggest that variation in MIRS associated with mosquito age and species can be successfully learned and traits predicted by a CNN from several laboratories, genetic backgrounds and environments across the three age groups (Fig. 4b). By using transfer learning, the ability to interpret the chemical signature of mosquito spectra (Fig. 2a) generated by the convolution layers of the model (dashed box Fig. 2e) is retained and can be adapted to learn the variation associated with ecologically diverse mosquitoes by utilising small quantities of these samples in the retraining. This demonstrates that transfer learning is an efficient and promising approach for minimising the amount of EV needed for recalibration and making DL-MIRS readily transportable to other mosquito populations.

### DL-MIRS detects age structure shifts in mosquito populations following vector control interventions

Next, we evaluated how well DL-MIRS could detect the impacts of vector control on wild mosquito populations. We have previously demonstrated that models trained on MIRS from laboratory-reared mosquitoes can be utilised to reconstruct the age structure of synthetic populations from which they were sampled and whether those populations had been subjected to LLINs^23^. However, it remains unclear if our present DL-MIRS model can achieve similar performance on genetically- and ecologically diverse mosquitoes. We therefore investigated whether DL-MIRS could detect changes in the age structure of simulated mosquito populations subjected to two common vector control interventions: the first has a rapid killing effect using long-lasting insecticide-treated nets (LLIN); the second primarily impacts “old” mosquitoes with attractive toxic sugar baits (ATSB) (Fig. 4b). To do this we first simulated the age structure of mosquito populations before and after LLIN or ATSB intervention assuming 36% mortality for mosquitoes above three days old or constant 9% mortality, respectively. Then, we estimated the statistical power of DL-MIRS to detect shifts in the three age groups (non-infected, potentially infected, potentially infectious) anticipated from these interventions (Fig. 4c,d, dotted line). We assessed how the power to detect age structure shifts varied with the number of local mosquitoes used for model testing (Supplementary Table S4). In both intervention scenarios, sampling 300 mosquitoes pre- and post-intervention was sufficient to obtain >80% power to detect an age structure shift when the training set was composed of 162 EV mosquito spectra (Fig. 4c,d, solid lines; and Supplementary Fig. 2). This shows that with relatively minor sampling and machine learning training efforts, this approach is capable of detecting population age structure shifts following vector control interventions.

## Discussion

Overall, our newly-developed DL-MIRS method aims to assist malaria mosquito surveillance by providing simultaneous information on mosquito species and age. The ability of DL-MIRS to discriminate mosquito age and species from MIRS and to generalised to new settings and populations by utilising a small proportion of the target population and transfer learning from our existing model, showing that the approach is flexible and scalable. The age determination for the training could be obtained either by rearing larvae from wild populations to adulthood in semi-field systems, or by morphological identification of the ovaries to determine gonotrophic cycles (e.g. Polovodova^7^). While both of these approaches are time consuming and require appropriate facilities, our results suggest that only a relatively small “one-off” sample size may be required for the re-training and the detection impacts of vector control interventions.

In the future, the accuracy of DL-MIRS for mosquito analyses could be substantially enhanced through enrichment of training sets with more spectra from additional populations and colonies. Additionally, inclusion of environmental data known to influence ageing rate, such as temperature, is likely to further increase prediction accuracy and generalisability. Consequently, DL-MIRS holds great promise and potential for integration into vector surveillance where it could play a key role in enhancing control and winning the fight against mosquito-borne diseases such as malaria.

## Methods

### Study sites

The experimental studies were conducted in two leading African malaria vector control institutions: Ifakara Health Institute (IHI), Tanzania and Institut de Recherche en Sciences de la Santé (IRSS), Burkina Faso. In IHI, the study experiments were conducted in the mosquito biology laboratory Vector Sphere and the larvae were collected from two villages in the Kilombero floodplains in Ulanga district, south-eastern Tanzania: Minepa village (longitude −8.285°, latitude 36.669°) and Tulizamoyo village (longitude −8.348°, latitude 36.732°). The ecology and species available in Ulanga district were described by Kaindoa et al^26^. In IRSS, mosquito sampling was conducted in the north of Bobo-Dioulasso in Vallée du Kou village (longitude −4.4201°, latitude 11.3824°) and in the south of Bobo-Dioulasso in Soumousso (longitude −4.0438°, and latitude 11.0125°).

### Mosquito rearing

We collected *Anopheles arabiensis, An. gambiae* and *An. coluzzii* mosquitoes born either from lab colonies or from wild mosquitoes, and reared either in the laboratory (at University of Glasgow [UoG], IHI or IRSS) or semi-field environments (at IHI or IRSS) (Fig. 1). Laboratory- and semi-field specific methods are described below.

#### Laboratory colonies

Mosquitoes were reared in the three different insectaries maintained under controlled temperature and humidity and a 12hrs:12hrs (light:dark) photoperiod, following standard operations as previously described^27^. Adult mosquitoes were fed with 5-10% sugar solution *ab libitum* via filter paper. Blood feeding was provided to allow egg production. Blood meals were provided using human blood at IHI directly by a human arm and at UoG through membrane feeding following^28^, and rabbit blood at IRSS directly on the animal. In each institution, different malaria vector species and strains were reared, as indicated in Supplementary Table 1. To produce age-matched mosquitoes, pupae were added to a clean cage on the same day. To generate different reproductive conditions, they were blood fed at different days after emergence and allowed to lay eggs in an oviposition cup 2 days after each blood meal. Mosquitoes were collected either 2 days after a blood meal (i.e. before egg laying) or 4 days after the blood meal (i.e. after egg laying had occurred). Mosquitoes were starved for 6-12h by removing sugar prior to blood feeding and each cage was blood fed every 6 days. Thus, mosquitoes living 6 or more days after their first blood meal underwent multiple gonotrophic cycles. Mosquitoes were sampled at ages ranging from 1 to 17 days old. A hundred and twenty mosquitoes per day (age) were assessed comprising each of the three physiological status.

#### Field-collected mosquitoes reared in the laboratory

At the Ifakara Health Institute, *An. arabiensis* larvae at different stages were collected from the study villages at different breeding habitats. Larvae were brought to the insectary and were sorted based on their morphology. The larvae were maintained in field water and provided with ground fish food (TetraMin^®^) until pupation. A plastic pipette was used to transfer pupae from the basins into disposable cups, which were then placed inside 30×30×30 cm cages until they emerged as adults. Emerged female and male adult mosquitoes were kept together to allow mating. These adult field mosquitoes were maintained at a same temperature 27±1.0°C, humidity 80±5% and a 12hrs:12hrs (light:dark) photoperiod, as lab-reared mosquitoes as previously described. At IRSS, all female mosquitoes, whether blood fed or gravid, were collected by trained technicians with mouth aspirators from local houses where mosquitoes have rested after a blood feeding. After aspiration the mosquitoes were transferred immediately into 30×30×30 cm cages covered with a wet cloth to avoid dehydration during transport. These mosquitoes were transferred to a room where light, humidity, and temperature are similar to that of the field (semi-field facility) and were maintained with glucose 5% for 72 hours to allow them to digest the blood. Then individual gravid females were transferred to a single cup containing 10ml water to allow oviposition. After two days, the females that laid eggs were removed with forceps and fixed in 80% ethanol for molecular species identification as previously described^29^. After the oviposition, only *An. gambiae* and *An. coluzzii* offspring from Soumousso and Vallée du Kou were kept and reared until adulthood for MIRS sample collection. At both sites, blood feeding and oviposition occurred in the same way and with the same timings as described for laboratory mosquitoes and a hundred and twenty mosquitoes per age (1 to 17 days) were assessed.

#### Laboratory colonies reared in the semi-field

At each site, experiments were conducted during the rainy season, which is the peak season for most mosquito species as it ensures optimal living conditions. Temperature and humidity were monitored every day in the semi-field and recorded. In addition, mosquitoes had access to live cattle for blood feeding and the semi-field chamber included water containers for mosquitoes to lay eggs in. These water containers were checked each day that adult collections were performed. The water was discarded and replaced on a daily basis in order to prevent the emergence of new adults. This ensured a single age group was present in each semi-field enclosure. Pupae from mosquito colonies (Supplementary Table 1) where released into the semi-field on two consecutive days and then recaptured at specific days for MIRS sampling. Day 0 was considered the day after the last batch of pupae was released into the facility. At IHI, mosquitoes were collected from Day 1 to 17 in batches of 100 mosquitoes per species and per age, while at IRSS mosquitoes were collected on Days 1, 4, 7, 10, and 15 in batches of 50 mosquitoes per species and per age.

### Spectroscopy

Upon collection of mosquitoes for all experiments conducted either in the laboratory, field or semi-field, mosquitoes were firstly transferred into a cup and then killed with cotton-soaked chloroform as previously described^23^ and then measured with MIRS. For each mosquito the average of >30 MIRS scans was used for subsequent analysis. Mosquito spectra were cleaned and minor atmospheric intrusion compensated, while those with low intensity or a significant atmospheric intrusion were discarded automatically using a custom script^30^ as previously described^23^.

### Laboratory and semi-field mosquito MIRS datasets

The core datasets used are detailed in Supplementary Table 2 and modifications of these datasets used for studies provided in the supplementary text are detailed in Table 3. All data sets are used for the prediction of mosquito age and species where age is provided as a categorical variable grouped into ages 1-4 days, 5-10 days, and 11+ days, and species is also categorical representing the given species of mosquito. These tables give details of the dataset that was used for each study, this includes number of data points, source of data, and what results supplied using these studies made us of as the testing set.

### Machine learning: building DL-MIRS

We trained a deep convolutional neural network (CNN) using 1-D convolutional layers to predict both age and species from MIRS input data. We chose CNNs because mid-infrared spectra are composed of multiple peaks which capture the biochemical characterization of mosquitoes; a network including numerous convolutional layers is capable of capturing complex local features in the spectra. In addition, fully connected layers are able to combine features learned by the convolutional layers to capture the correlation of features across the entire spectra, a necessity for the high dimensional spectra used for analysing mosquito cuticles. Each 1-D convolutional layer defines filters of fixed width with trainable weights and biases, where each convolutional operation is the sum of the dot product between the filter and the section of the spectra currently considered. The assumption of locality made in convolutional neural networks holds when using mid-infrared spectra due to fixed width wavenumber bands corresponding to individual vibrational modes. Therefore, convolutional layers are able to learn local structure in the spectra. In addition, we applied batch normalisation to improve the stability of the neural network and max pooling was used to reduce the spatial size of the representation. Further, *L*_2_ regularisation was used in each layer to reduce overfitting, with a convolutional stride of size one in convolutional layers one, three and five, and stride of two in convolutional layers two and three. We used size two max pooling in the final convolutional layer and dropout was applied before the dense layer to further reduced overfitting. The convolutional neural network architecture was found by optimising the hyperparameters. For this, the number of layers was hand optimised, whilst the kernel, stride, and pooling sizes for each convolutional layer were optimised using gp_minimize from the scikit-optimize package^31^.

Unless otherwise stated, we trained the DL-MIRS CNN using datasets balanced across mosquito age groups and species. We began by splitting out 10% of the dataset stratified by age and species for subsequent testing of the trained models. The remaining 90% of the dataset was used for optimising models through 10-fold cross validation. Both age and species groups were binarised using MultiLabelBinarizer^31^ and the spectra were standardised using StandardScaler^31^ to centre each variable around its global mean and scale it to unit variance. Machine learning was performed in Python 3.6.8 using keras 2.2.4^32^ and tensorflow-gpu 1.12.0^33^.

### Power estimation of vector control detection

To estimate the statistical power of the DL-MIRS model to detect a shift in the age structure of a mosquito population after each of two common insecticidal interventions, we generated computer simulated mosquito populations exposed to long-lasting insecticide-treated nets (LLIN), and of attractive toxic sugar baits (ATSB). The age structure (i.e., the frequency of each age class) in the control (pre-intervention) population was simulated assuming a constant daily mortality of 9%^34^ up to 20 days, with no survival after 20 days. The LLIN intervention was assumed to cause a death rate of 36%, four times higher than natural mortality, but applying only after day 3 assuming mosquitoes will host seek and encounter a bed-net only after this age. In the population exposed to the ATSB intervention, mortality was assumed to be two times higher (18%) than natural mortality, applying throughout the mosquitoes’ lives. The age structure of each post-intervention population was then compared with the control population using Wilcoxon/Mann-Whitney U tests. Power was estimated across all 70 combinations of seven sample sizes (n = 20, 50, 100, 150, 200, 250, and 300) and ten levels of enrichment of the training data with EV data (0–17% of the training data was EV data). For each of these 70 scenarios, power was estimated as the proportion of 10,000 simulated data sets where a significant (P < 0.05) difference in age structure was detected between intervention and control populations. Simulations were performed in R version 3.6.1^35^.

### Ethical statement

The present study has been agreed by the institutional ethical committee of Institut de Recherche en Sciences de la Santé (IRSS) under the number A012-2017/CEIRES on 03 July 2017 before its implantation on the sites. At Ifakara Health Institute, Ethical approval for the study was obtained from the Ifakara Health Institute Institutional Review Board (Ref. IHI/IRB/EXT/No: 005-2018), and from the Medical Research Coordinating Committee (MRCC) at the National Institutes of Medical Research (NIMR), Ref: NIMR/HQ/R.8c/Vol.II/880. At the University of Glasgow, human blood for feeding female mosquitoes was obtained from the Glasgow and West of Scotland Blood Transfusion Service. Ethical approval for the supply and use of human blood was obtained from Scottish National Blood Transfusion Service committee for governance of blood and tissue samples for non-therapeutic use, and Donor Research (submission Reference No 18 15). Whole blood from donors of any blood group was provided in Citrate-Phosphate-Dextrose-Adenine (CPD-A) anticoagulant/preservative. Fresh blood was obtained on a weekly basis.

### Data & code availability

All relevant data are available from the corresponding authors upon reasonable request. All code used for machine learning and power analysis will be made available upon peer-reviewed publication.

## Acknowledgements

This work was funded by the Medical Research Council GCRF Infections Foundation Awards MR/P025501/1. SAB, FB, and FO are supported by the Royal Society International Collaboration Award ICA/R1/191238 and Bill and Melinda Gates Foundation award OPP1217647. FO was also supported by a Wellcome Trust Intermediate Fellowship in Public Health and Tropical Medicine (Grant Number: WT102350/Z/13), FB by an AXA RF fellowship (14-AXA-PDOC-130) and an EMBO LT fellowship (43-2014). KW and MGJ thank the Engineering and Physical Sciences Research Council (EPSRC) for support through grants EP/K034995/1, EP/N508792/1, and EP/N007417/1, the Leverhulme Trust through Research Project Grant RPG-2018-350, and the European Research Council (ERC) under the European Union’s Horizon 2020 research and innovation program (grant agreement No. 832703). JM is supported by a University of Glasgow Lord Kelvin Adam Smith Studentship. RM-S is grateful for EPSRC support through grants EP/R018634/1 and EP/T00097X/1. We would like to thank Dorothy Armstrong and Elizabeth Peat for assistance with mosquito rearing and maintenance. We would also like to thank Hilary Ranson for providing the Kisumu colony. We thank Fibios Science Communication for help with graphics.

## Author contributions

Conceptualisation: DJS, RS, JM, KW, RM-S, HF, MG-J, SAB, AD, FOO, FB. Data curation: DJS, RS, JM, EPM, AN, IS, MG-J, FB. Formal Analysis: JM, PCDJ, RM-S, MG-J, SAB. Funding acquisition: KW, RM-S, HF, AD, FOO, FB. Investigation: DJS, RS, EPM, AN, IS, MG-J, FB. Methodology: DJS, RS, JM, EPM, AN, PCDJ, GF, KW, HF, MG-J, SAB, AD, FOO, FB. Project administration: DJS, RS, AN, HF, AD, FOO, FB. Resources: AMGB, KW, HF, AD, FOO, FB. Software: JM, RM-S, MG-J, SAB. Supervision: AMGB, KW, RM-J, HF, MG-J, SAB, AD, FOO, FB. Validation: JM, PCDJ, RM-S, MG-J, MG-J, SAB, FB. Visualization: DJS, RS, JM, PCDJ, MG-J, SAB, FB. Writing – original draft: DJS, RS, JM, PCDJ, MG-J, SAB, FB. Writing – review & editing: all authors.

## References

1. Beier, J. Malaria parasite development in mosquitoes. Annu Rev Entomol 43, 519–543 (1998).

2. MacDonald, G. Epidemiological basis of malaria control. Bull World Health Organ 15, 613–626 (1956).

3. MacDonald, G. II. The objectives of residual insecticide campaigns. Transactions of the Royal Society of Tropical Medicine and Hygiene 46, 227–235 (1952).

4. Bhatt, S. et al. The effect of malaria control on *Plasmodium falciparum* in Africa between 2000 and 2015. Nature 526, 207–211 (2015).

5. Churcher, T., Lissenden, N., Griffin, J., Worrall, E. & Ranson, H. The impact of pyrethroid resistance on the efficacy and effectiveness of bednets for malaria control in Africa. Elife 5 (2016).

6. Beklemishev, W., Detinova, T. & Polovodova, V. Determination of physiological age in anophelines and of age distribution in anopheline populations in the USSR. Bull World Health Organ 21, 223–232 (1959).

7. Hugo, L. E., Quick-miles, S., Kay, B. H. & Ryan, P. A. Evaluations of Mosquito Age Grading Techniques Based on Morphological Changes. Journal of Medical Entomology 45, 353–369 (2008).

8. Johnson, B., Hugo, L., Churcher, T., Ong, O. & Devine, G. Mosquito age grading and vector-control programmes. Trends Parasitol (2019).

9. Schlein, Y. Age grouping of anopheline malaria vectors (Diptera: Culicidae) by the cuticular growth lines. Journal of Medical Entomology 16, 502–506 (1979).

10. Caputo, B. et al. Identification and composition of cuticular hydrocarbons of the major Afrotropical malaria vector *Anopheles gambiae s.s*. (Diptera: Culicidae): analysis of sexual dimorphism and age-related changes. J Mass Spectrom 40, 1595–1604 (2005).

11. Cook, P. et al. Predicting the age of mosquitoes using transcriptional profiles. Nat Protoc 2, 2796–2806 (2007).

12. Bass, C., Williamson, M., Wilding, C., Donnelly, M. & Field, L. Identification of the main malaria vectors in the *Anopheles gambiae* species complex using a TaqMan real-time PCR assay. Malar J 6, 155 (2007).

13. Mayagaya, V. et al. Non-destructive determination of age and species of *Anopheles gambiae s.l*. using near-infrared spectroscopy. Am J Trop Med Hyg 81, 622–630 (2009).

14. Sikulu, M. et al. Mass spectrometry identification of age-associated proteins from the malaria mosquitoes *Anopheles gambiae s.s*. and *Anopheles stephensi*. Data Brief 4, 461–467 (2015).

15. Ferguson, H. et al. Ecology: a prerequisite for malaria elimination and eradication. PLoS Med 7, e1000303 (2010).

16. Cohuet, A., Harris, C., Robert, V. & Fontenille, D. Evolutionary forces on Anopheles: what makes a malaria vector. Trends Parasitol 26, 130–136 (2010).

17. Ferguson, H. et al. Establishment of a large semi-field system for experimental study of African malaria vector ecology and control in Tanzania. Malar J 7, 158 (2008).

18. Lambert, B. et al. Monitoring the Age of Mosquito Populations Using Near-Infrared Spectroscopy. Sci Rep 8, 5274 (2018).

19. Peiris, K. H., Drolet, B. S., Cohnstaedt, L. W. & Dowell, F. E. Infrared absorption characteristics of culicoides sonorensis in relation to insect age. American Journal of Agricultural Science and Technology (2014).

20. Krajacich, B. et al. Analysis of near infrared spectra for age-grading of wild populations of *Anopheles gambiae*. Parasit Vectors 10, 552 (2017).

21. Waynant, R. W., Ilev, I. K. & Gannot, I. Mid–infrared laser applications in medicine and biology. Philosophical Transactions of the Royal Society of London. Series A: Mathematical, Physical and Engineering Sciences 359, 635–644 (2001).

22. Sorak, D. et al. New developments and applications of handheld raman, mid-infrared, and near-infrared spectrometers. Applied Spectroscopy Reviews 47, 83–115 (2012).

23. González Jiménez, M. et al. Prediction of mosquito species and population age structure using mid-infrared spectroscopy and supervised machine learning. Wellcome Open Res 4, 76 (2019).

24. Sigurdsson, S. et al. Detection of skin cancer by classification of Raman spectra. IEEE Trans Biomed Eng 51, 1784–1793 (2004).

25. McInnes, L., Healy, J. & Melville, J. UMAP: Uniform manifold approximation and projection for dimension reduction. arXiv, 1802.03426v2 (2018).

26. Kaindoa, E. et al. Housing gaps, mosquitoes and public viewpoints: a mixed methods assessment of relationships between house characteristics, malaria vector biting risk and community perspectives in rural Tanzania. Malar J 17, 298 (2018).

27. Siria, D. et al. Evaluation of a simple polytetrafluoroethylene (PTFE)-based membrane for blood-feeding of malaria and dengue fever vectors in the laboratory. Parasit Vectors 11, 236 (2018).

28. Carter, R., Ranford-Cartwright, L. & Alano, P. The culture and preparation of gametocytes of *Plasmodium falciparum* for immunochemical, molecular, and mosquito infectivity studies. Methods Mol Biol 21, 67–88 (1993).

29. Santolamazza, F. et al. Insertion polymorphisms of SINE200 retrotransposons within speciation islands of *Anopheles gambiae* molecular forms. Malar J 7, 163 (2008).

30. Gonzalez Jimenez, M. https://github.com/SimonAB/Gonzalez-Jimenez_MIRS/blob/master/Locomosquito.ipynb (2019).

31. Pedregosa, F. et al. Scikit-learn: Machine Learning in Python. Journal of Machine Learning Research 12, 2825–2830 (2011).

32. Chollet, F. et al. Keras https://keras.io. 2015.

33. Pasumarthi, R. K. et al. TF-Ranking: Scalable TensorFlow Library for Learning-to-Rank in Proceedings of the 25th ACM SIGKDD International Conference on Knowledge Discovery and Data Mining (Anchorage, AK, 2019), 2970–2978.

34. Molineaux, L., Gramiccia, G. & World Health Organization. The Garki project: research on the epidemiology and control of malaria in the Sudan savanna of West Africa (World Health Organization, 1980).

35. R Core Team. R: A language and environment for statistical computing (2019).

36. MR4. BEI Resources Knowledge Base, https://www.beiresources.org.

37. Nsango, S. E. et al. Genetic clonality of *Plasmodium falciparum* affects the outcome of infection in Anopheles gambiae. Int. J. Parasitol. 42, 589–595 (2012).

38. Lyimo, I. et al. The impact of host species and vector control measures on the fitness of African malaria vectors. Proc Biol Sci 280, 20122823 (2013).

39. Ng’habi, K. et al. Effect of larval crowding on mating competitiveness of *Anopheles gambiae* mosquitoes. Malar J 4, 49 (2005).

40. Ng’habi, K., Mwasheshi, D., Knols, B. & Ferguson, H. Establishment of a self-propagating population of the African malaria vector *Anopheles arabiensis* under semi-field conditions. Malar J 9, 356 (2010).

41. Bilgo, E. et al. Transgenic *Metarhizium pingshaense* synergistically ameliorates pyrethroid-resistance in wild-caught, malaria-vector mosquitoes. PLoS One 13, e0203529 (2018).

