## Supplementary Information for "Rapid ageing and species identification of natural mosquitoes for malaria surveillance"

### Supplementary Tables

| Location | Species | Strain | Reference |
|---|---|---|---|
| Glasgow, UK | *An. gambiae* | Kisumu | [36] |
| | *An. coluzzii* | Ngousso | [37] |
| | *An. arabiensis* | Ifakara | [38] |
| Ifakara, TZ | *An. gambiae* | Ifakara, Niage | [39] |
| | *An. arabiensis* | Ifakara, Niage | [40] |
| Bobo, BF | *An. gambiae* | Soumousso | [41] |
| | *An. coluzzii* | Vallée du Kou | [41] |

**Supplementary Table 1** List of species and strains used for training DL-MIRS.

| **Study Laboratory Variation (LV)** |
|---|
| Data from group LV only. |
| 8224 mosquito data points. |
| Data from Tanzania, Burkina Faso, United Kingdom. |
| Data balanced, where possible, by country, species, and age groups. |
| **Study Genetic Variation (GV)** |
| Data from groups LV and GV. |
| 4800 mosquito data points from group LV and 2400 mosquito data points from group GV. |
| Data from Tanzania and Burkina Faso only. |
| Data balanced by country, species, and age groups. |
| Testing data set is a 10% split of group GV data only. |
| **Study Environmental Variation (EV)** |
| Implicit use of Data from groups LV and GV. |
| Data from group EV. |
| A varying number of data points from the EV group. |
| EV data from Tanzania and Burkina Faso only. |
| Data balanced by country, species, and age groups for groups LV and GV, and balanced where possible for group EV. |
| Testing data set is a hold out data set of 180 EV data points not included in the training set. |

**Supplementary Table 2** Allocation of samples between training and testing of DL-MIRS

| **E1 – Reducting parameters** |
|---|
| Data from groups LV, GV, and EV only. |
| 4800 mosquito data points from group LV and 2400 mosquito data points from group GV in the training set. |
| Data from Tanzania and Burkina Faso only. |
| Data balanced by country, species, and age groups for groups LV and GV. |
| Testing data set is group EV. |
| Consider how the number of trainable parameters in the model affects classification accuracy. |
| **E2 – Cross EV** |
| Data from groups GV, and EV. |
| 2400 mosquito data points from group GV in the training set and either 1200 data points from Tanzania or 300 data points from Burkina Faso from Group EV. |
| Data from Tanzania and Burkina Faso only. |
| Data balanced by country, species, and age groups for groups GV. Data balanced by species, and age groups for groups EV. |
| Testing data set is data from the opposite country to that which was included during training from group EV. |
| Attempt to classify unseen EV data from another country. |
| **E3 - Cross laboratory** |
| Data from group LV only. |
| 8224 mosquito data points. |
| Data balanced, where possible, by country, species, and age groups. |
| Leave data out from one country as the testing set in an attempt to classify data from an unseen lab. |
| **E4 – Cross laboratory with reduced parameters** |
| Data from group LV only. |
| *Stage 1:* |
| Initial training set is 3424 mosquito data points from UK group LV, balanced where possible by species, and age groups. |
| Sensitivity analysis performed on model from stage 1 to select frequencies. |
| *Stage 2:* |
| Training set is 2400 mosquito data points from either Tanzania or Burkina Faso in group LV, balanced by species, and age groups. |
| Testing data set is from group LV from the country left out during training. |

**Supplementary Table 3** Grouping of mosquito subsets for cross-validation and generalisation of DL-MIRS.

**Supplementary Table 4** Predicted power to detect a shift in age structure in response to each of two interventions, long-lasting insecticide-treated nets (LLIN) and attractive toxic sugar baits (ATSB) relative to a population with no intervention (see Fig. 4). Power depended on the number of mosquitoes sampled from each population (intervention and control) and the number of spectra from semi-field (EV) mosquitoes used in the training set. Each power value was estimated from analysis of 10,000 simulated data sets.

| N sampled | N EV spectra added | Power | |
|---|---|---|---|
| | | LLIN intervention | ATSB intervention |
| 20 | 0 | 6.9% | 5.0% |
| 20 | 162 | 46.1% | 14.8% |
| 20 | 324 | 60.0% | 18.6% |
| 20 | 486 | 60.9% | 18.8% |
| 20 | 654 | 74.6% | 22.7% |
| 20 | 815 | 78.0% | 24.4% |
| 20 | 973 | 75.0% | 23.4% |
| 20 | 1131 | 79.0% | 24.3% |
| 20 | 1294 | 80.7% | 25.8% |
| 20 | 1452 | 78.5% | 24.9% |
| 50 | 0 | 10.3% | 6.4% |
| 50 | 162 | 84.6% | 31.1% |
| 50 | 324 | 94.9% | 40.5% |
| 50 | 486 | 95.2% | 41.7% |
| 50 | 654 | 98.8% | 50.5% |
| 50 | 815 | 99.3% | 53.8% |
| 50 | 973 | 98.9% | 51.2% |
| 50 | 1131 | 99.4% | 54.3% |
| 50 | 1294 | 99.4% | 55.2% |
| 50 | 1452 | 99.2% | 54.4% |
| 100 | 0 | 15.4% | 7.5% |
| 100 | 162 | 99.1% | 55.8% |
| 100 | 324 | 99.9% | 69.5% |
| 100 | 486 | 99.9% | 70.0% |
| 100 | 654 | 100.0% | 80.5% |
| 100 | 815 | 100.0% | 83.2% |
| 100 | 973 | 100.0% | 80.8% |
| 100 | 1131 | 100.0% | 82.9% |
| 100 | 1294 | 100.0% | 84.7% |
| 100 | 1452 | 100.0% | 83.2% |
| 150 | 0 | 20.6% | 8.7% |
| 150 | 162 | 99.9% | 73.5% |
| 150 | 324 | 100.0% | 85.6% |
| 150 | 486 | 100.0% | 86.4% |
| 150 | 654 | 100.0% | 93.6% |
| 150 | 815 | 100.0% | 94.7% |
| 150 | 973 | 100.0% | 93.3% |
| 150 | 1131 | 100.0% | 94.7% |
| 150 | 1294 | 100.0% | 95.4% |
| 150 | 1452 | 100.0% | 95.4% |
| 200 | 0 | 25.9% | 10.6% |
| 200 | 162 | 100.0% | 84.1% |

*Continued on next page...*

**Table 4 continued from previous page**

| N sampled | N EV spectra added | Power | |
|---|---|---|---|
| | | LLIN intervention | ATSB intervention |
| 200 | 324 | 100.0% | 93.9% |
| 200 | 486 | 100.0% | 94.6% |
| 200 | 654 | 100.0% | 97.8% |
| 200 | 815 | 100.0% | 98.7% |
| 200 | 973 | 100.0% | 98.2% |
| 200 | 1131 | 100.0% | 98.3% |
| 200 | 1294 | 100.0% | 98.7% |
| 200 | 1452 | 100.0% | 98.6% |
| 250 | 0 | 30.9% | 11.6% |
| 250 | 162 | 100.0% | 91.9% |
| 250 | 324 | 100.0% | 97.6% |
| 250 | 486 | 100.0% | 97.3% |
| 250 | 654 | 100.0% | 99.4% |
| 250 | 815 | 100.0% | 99.7% |
| 250 | 973 | 100.0% | 99.5% |
| 250 | 1131 | 100.0% | 99.6% |
| 250 | 1294 | 100.0% | 99.7% |
| 250 | 1452 | 100.0% | 99.6% |
| 300 | 0 | 36.5% | 13.4% |
| 300 | 162 | 100.0% | 95.3% |
| 300 | 324 | 100.0% | 99.1% |
| 300 | 486 | 100.0% | 99.2% |
| 300 | 654 | 100.0% | 99.7% |
| 300 | 815 | 100.0% | 100.0% |
| 300 | 973 | 100.0% | 99.8% |
| 300 | 1131 | 100.0% | 100.0% |
| 300 | 1294 | 100.0% | 100.0% |
| 300 | 1452 | 100.0% | 99.9% |

505 ## Supplementary Figures

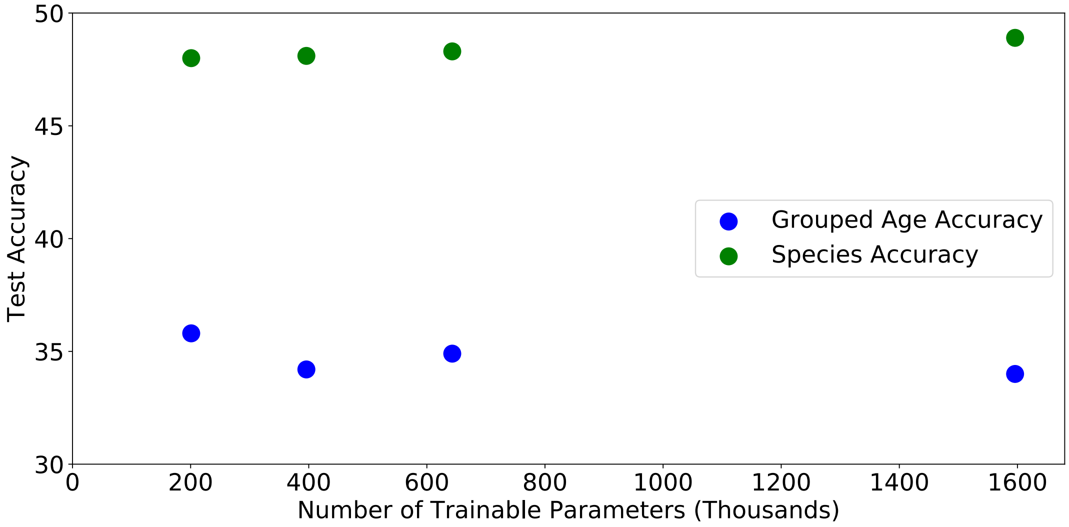


**Supplementary Fig. 1 Model overfitting.** The CNN was modified to reduce capacity, lowering the number of trainable parameters, to explore the effect on classification accuracy with a testing dataset comprising of EV data only, while the training dataset comprises of LV and GV data. This suggests that the model is not overfitting in the case of training on LV and GV data only, as the model does not increase testing accuracy on EV data when lowering the number of trainable parameters.

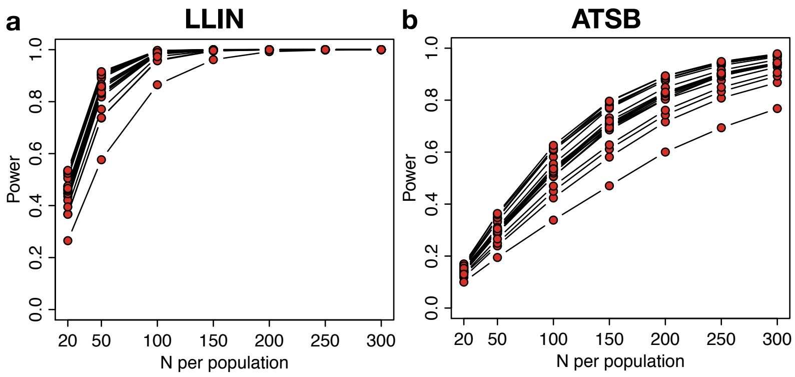


**Supplementary Fig. 2** Impact of the number of mosquitoes used for DL-MIRS training on power to detect an effect of vector control on mosquito population age structure for each of two vector control interventions, **a**, long-lasting insecticide-treated nets (LLIN) and **b**, attractive toxic sugar baits (ATSB) relative to a population with no intervention (control).

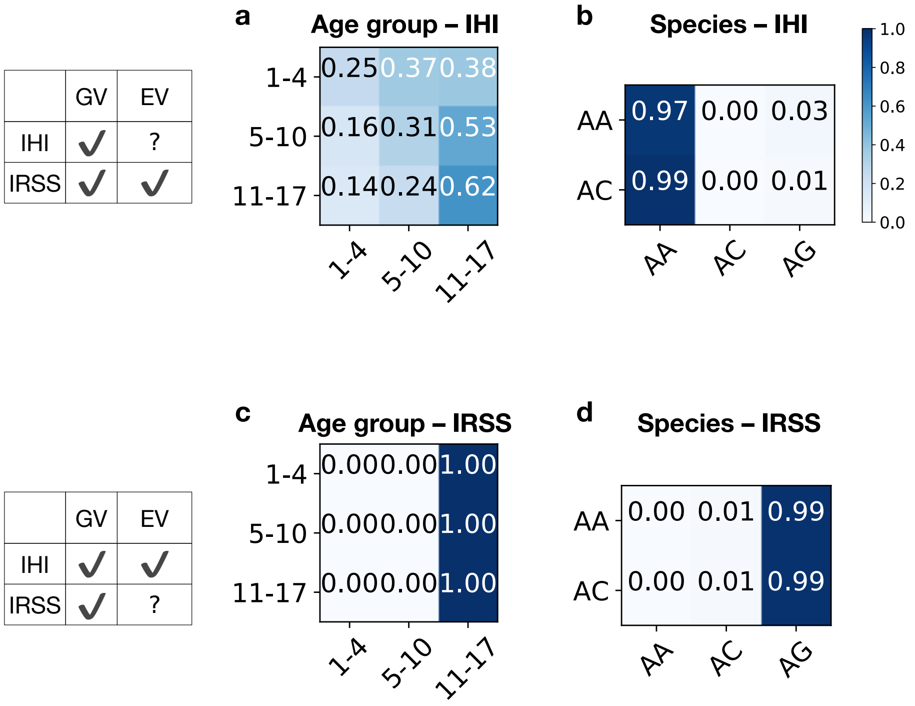


**Supplementary Fig. 3 EV data from an unseen site.** All of the EV samples from one site were held out and DL-MIRS was trained on the rest of GV and EV samples from IHI and IRSS. This demonstrates that if the training dataset only has GV data for a site it will not be possible for the model to classify EV data, despite the inclusion of EV data from other sites.

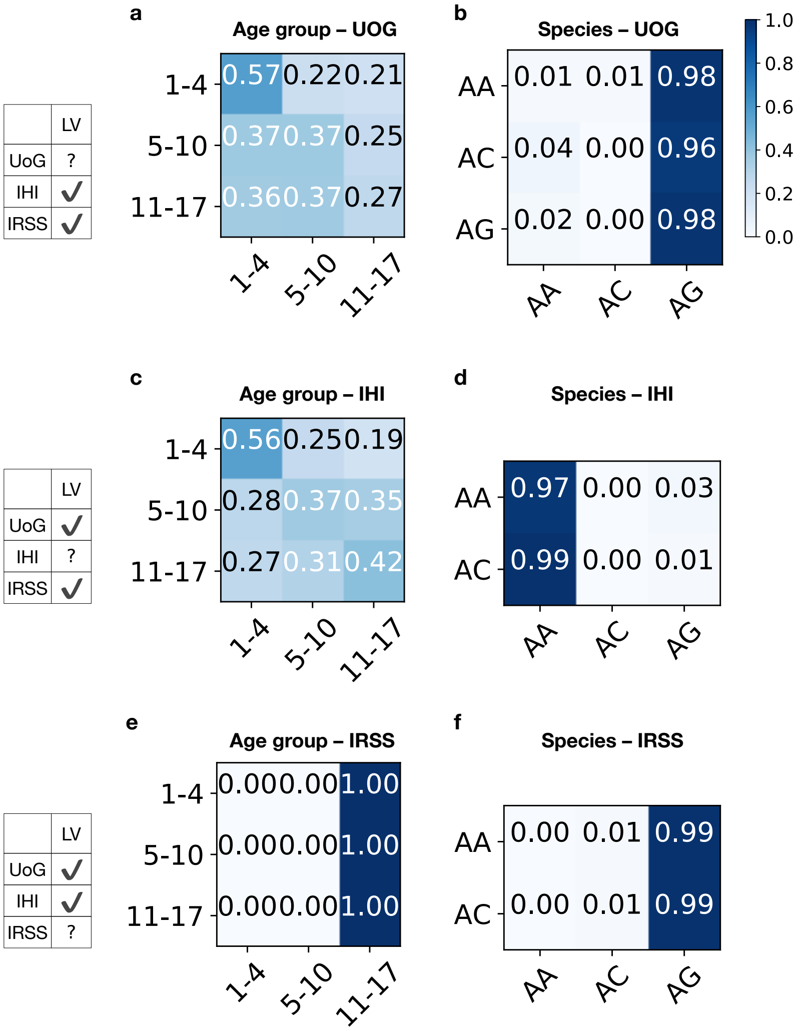


**Supplementary Fig. 4 LV data from an unseen site.** DL-MIRS was trained with LV data from two sites and then used to classify LV data from a third site. The results demonstrate that the model is not able to learn features capable of classifying data from a previously unseen laboratory. This shows that differences exist in the MIRS data across laboratories from different sites. The result is in agreement with the initial UMAP data exploration, demonstrating differences across laboratories.

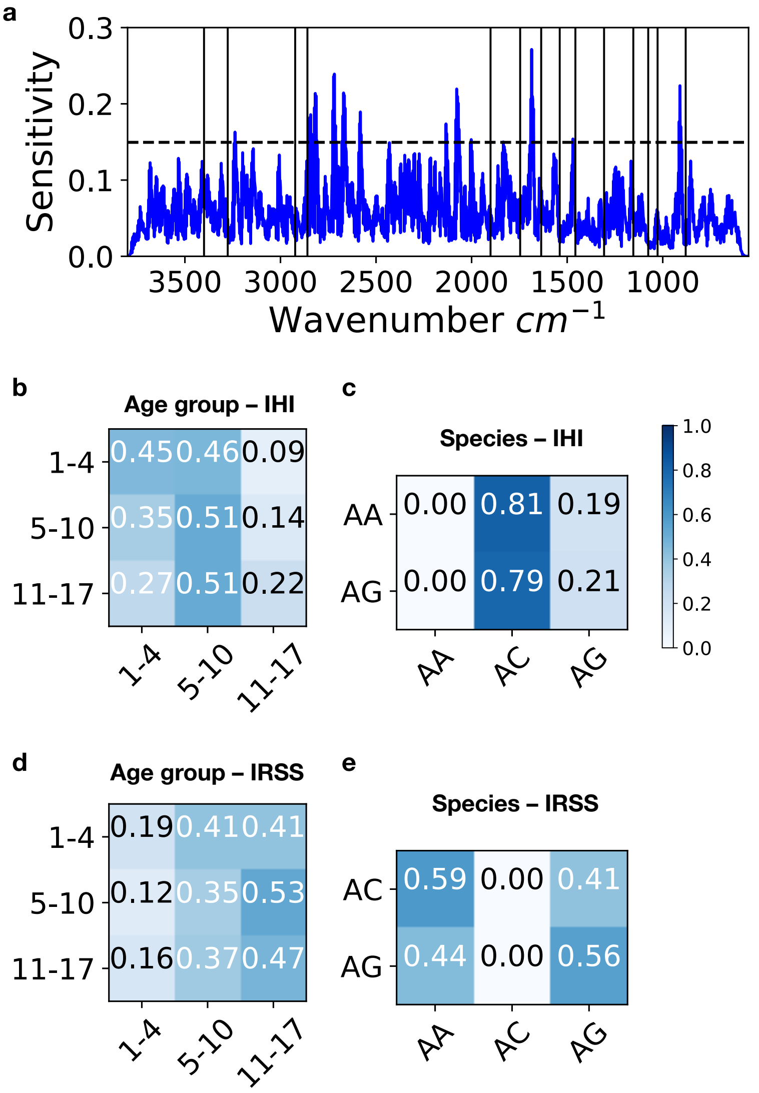


**Supplementary Fig. 5 Preselecting wavenumbers from sensitivity analysis of a CNN.** The sensitivity plot shows the model sensitivity to differing wavenumbers for a model trained with LV data from the UK only, with the dashed blue line representing a 95% confidence in the sensitivity values[24]. A fully connected neural network was then trained, where the input was the wavenumbers identified from the sensitivity analysis on UK data, on IHI and IRSS LV data separately. These models were then tested on LV data from either IHI or IRSS depending which was left out during training. These results demonstrate that no performance increase can be gained by preselecting wavenumbers, further aiding the argument that the deep CNN model is not over fitting when generalisation is not demonstrated.
